# Exposure to environmental stress decreases the resistance of river microbial communities towards invasion by antimicrobial resistant bacteria

**DOI:** 10.1101/2022.11.19.517188

**Authors:** Kenyum Bagra, Xavier Bellanger, Christophe Merlin, Gargi Singh, Thomas U. Berendonk, Uli Klümper

## Abstract

Environmental microbiomes are constantly exposed to invasion events through foreign, antibiotic resistant bacteria that were enriched in the anthropic sphere. However, the biotic and abiotic factors, as well as the natural barriers that determine the invasion success of these invader bacteria into the environmental microbiomes are poorly understood. A great example of such invasion events are river microbial communities constantly exposed to resistant bacteria originating from wastewater effluents. Here, we aim at gaining comprehensive insights into the key factors that determine their invasion success with a particular focus on the effects of environmental stressors, regularly co-released in wastewater effluents. Understanding invasion dynamics of resistant bacteria is crucial for limiting the environmental spread of antibiotic resistance. To achieve this, we grew natural microbial biofilms on glass slides in rivers for one month. The biofilms were then transferred to laboratory, recirculating flume systems and exposed to a single pulse of a model resistant invader bacterium (*E. coli*) either in presence or absence of stress induced by Cu^2+^. The invasion dynamics of *E. coli* into the biofilms were then monitored for 14 days. Despite an initially successful introduction of *E. coli* into the biofilms, independent of the imposed stress, over time the invader perished in absence of stress. However, under stress c the invading strain successfully established and proliferated in the biofilms. Noteworthy, the increased establishment success of the invader coincided with a loss in microbial community diversity under stress conditions, likely due to additional niche space becoming available for the invader.

## Introduction

The acquisition of resistance against antibiotics by human pathogens has become a global threat to human health. In a worst case scenario, it is predicted to claim up to 10 million lives by 2050 [1]. Within the context of the spread of antimicrobial resistance (AMR) it has been pointed out that applying a One Health approach is of high importance [2]. This consists of the basic idea that the health of humans, animals and the environment are interconnected. Consequently, it is vital to consider the surrounding environments in order to maintain human health [3].

One of the most evident reasons for the rise of AMR within environmental microbiomes [4–6] is the environmental contamination with resistant bacteria and, in some instances, selective as well as co-selective agents due to anthropogenic activities [7–9]. Water bodies, especially rivers, have, among others, been considered as one of the “hotspots” of environmental AMR pollution [10, 11], as many urban rivers receive a daily influx of wastewater effluents from various sources. These introduce foreign microbes (e.g. pathogenic and commensal bacteria from the human gut microbiome) which are regularly enriched in antibiotic resistant genes (ARGs) [12–14]. In addition, abiotic pollutants, such as pesticides [15, 16], pharmaceutical products [17, 18], personal care products [19, 20] and metal trace elements [7, 9, 19] are regularly co-introduced with the bacterial contaminants. These are known to exert selection pressures on ARB and furthermore lead to increased horizontal gene transfer rates which result in the amplification of ARGs once they entered the river microbiome [12, 20–23], hence making rivers a reservoir and breeding ground for AMR [5, 24, 25]. However, there is no clear understanding whether the co-introduction of stressors through, e.g., wastewater effluents, also affects the invasion success of the constantly incoming resistant bacteria into the river microbiomes.

A successful invasion by foreign bacteria into a microbial community proceeds in three main steps: a) introduction, b) establishment, and c) growth and spread, and d) impact [26]. The discharge of wastewater into the river introduces the foreign bacteria to a new environment. Thereafter, the invading bacteria have to survive not only the “abiotic” stresses in the new environment, but also to face potential intrinsic resistances from the indigenous microbial community, hence a biotic barrier towards invasion, to establish themselves. The invading bacteria have to overcome this biotic barrier by competing for niches and resources to persist and thrive. In this step, the invading bacteria can either succeed or fail to outcompete the indigenous bacteria [26–28]. During this process the availability of environmental niches for the invader is a crucial parameter. In general, long-term established, rich and diverse microbial communities occupy a wider set of niches in their given environment. Hence, there is less niche spaces available for foreign bacteria which limits their invasion success [26, 28–30]. Consequently, such long-term established native communities tend to usually be highly resistant to invasion. However, studies on rivers have demonstrated that the microbial community composition downstream of wastewater treatment plants, where a majority of invasion events may take place, is regularly altered due to the consistent inflow of environmental stressors [31–33]. Understanding the invasion dynamics of resistant bacteria into the river microbiome in the presence of environmental stress can help mitigate the spread of AMR by identifying the key factors that determine the naturally occurring barrier effects towards invasion. Invasion theory suggests that if a microbial community’s composition is disrupted, additional niche space becomes available resulting in a greater chance of the environment being invaded [29]. Within this theoretical framework, we hypothesized that, in the context of wastewater effluents, the success rate of invasion and the establishment of foreign resistant bacteria into the resident river microbial communities is higher when the community diversity and structure is disturbed due to the co-release of environmental stressors.

To test this hypothesis, we collected natural microbial biofilms of differing diversity and composition grown in two different pristine German rivers. These were transferred to a laboratory flume system and exposed to a model invading bacterium, here the resistant *E. coli* strain CM2372, in the presence and absence of stress induced by copper (Cu^2+^). *E. coli* was chosen as the model invader bacterium, as it is a usual fecal contaminant commonly used as an indicator of anthropogenic pollution through wastewaters [4, 6, 34–36]. Copper was selected as model stressor, due to its ubiquitous presence as a river contaminant [19, 21, 37] and its previous identification as a dominant stressor in causing (co-)selection and increased horizontal gene transfer of ARGs [21, 22, 38]. Invasion experiments were carried out for two weeks with biofilms destructively sampled in regular time intervals to determine the invasion success of *E. coli* into the biofilm in the presence and absence of copper induced stress.

## Material & Methodology

### *E. coli* model invader strain

*E. coli strain* CM2372 was used as the model invader bacterium. This strain derived from *E. coli* EDCM367 [39], an MG1655 derivative with a Δ*lacZY* deletion authorizing a specific qPCR primer design for the quantification of the bacterium. Strain CM2372 is differentiated from EDCM367 through the introduction of the broad host range IncP1-α plasmid pG527 encoding resistance to kanamycin by plate mating using the donor strain *Pseudomonas putida* UWC1::pG527 [40].

To test, whether the selected model stressor copper inhibited the model *E. coli* invader strain, its maximum growth rate in LB medium at 0 ppb, 250 ppb and 500 ppb CuSO_4_ was assessed spectrophotometrically at 600nm. Copper did not inhibit the growth of *E. coli* strain CM2372 at either 250ppb (µ_max_=0.017±0.005) or 500ppb (µ_max_=0.017±0.005) compared to the control (µ_max_=0.014±0.005) (all p>0.05, t-test, n=15), hence the highest tested concentration of 500 ppb (µg/L) was used for the subsequent copper treatment. Importantly, copper did not provide a selective advantage for any of the plasmid encoded resistance traits.

### River microbial community and river water

Natural river biofilm microbial communities were obtained from two rivers - Hausdorfer Bach (50°54’49.8”N, 13°47’13.9”E) and Hirschbach (50°54’17.6”N, 13°45’06.3”E) - both part of the Lockwitzbach tributary in Saxony, Germany. The chosen locations were picked to have no upstream influence of any wastewater treatment plant effluents. Biofilms were grown on microscope glass slides of 76 × 26 mm size (DWK Life science, Wertheim, Germany). Artificial Exposure Units (AEUs) constructed from Plexiglas (Fig. SI1) containing 9 glass slides each were screwed onto a concrete plate and covered with a stainless steel cover providing shade (limitation of photosynthetic organisms) and protection. A metal mesh at the front and the back allowed river water to stream through the AEUs. The AEUs were immerged in the rivers for one month in September 2021 to allow biofilm growth. The AEUs containing the glass slides with grown biofilms were then collected and individually transferred to the laboratory in a 1L box containing river water of the respective side at ambient temperature. Simultaneously, 100L of river water from each respective river were collected and filter sterilized using 0.2 µm pore size membrane filters (Whatman, Maidstone, UK).

### Laboratory flume system

Laboratory artificial flume systems were 40 cm long, 19.1 cm width and 10 cm height, made of polypropylene, and equipped with an in- and an outflow pipe at either end (Fig. SI1). Each flume was connected to a 2L reservoir connected to a recirculation pump and filled with a total volume of 6.348 L of filter sterilized river water constantly recirculated with a pump at 100ml/min. At one end, the inflow pipe (1 cm in diameter) was located at 0.5 cm height with the inflow stream directed toward the bottom of the flume to allow for an even distribution of flow. At the other end an overflow pipe was positioned centrally at 7 cm height in order to maintain a constant maximum water level inside the flume. Three replicate AEUs containing glass slides covered with river biofilms were placed in the middle of each flume filled with filter sterilized river water of the corresponding site.

### Treatments

Before starting the treatment process, the biofilms were set to acclimatize to the laboratory conditions for a week. The temperature was maintained at 20°C and flumes were kept in the dark throughout the experiment. After the acclimatization period, the flumes for each river, Hausdorfer Bach (HAU) and Hirschbach (HIR), were subject to three different types of treatments for a period of 14 days: a) solely exposed to *E. coli* CM2372 (-Cu), b) exposed to *E. coli* CM2372 and Cu^2+^ (+Cu) and c) a control group was run without exposure to either *E. coli* CM2372 or Cu^2+^. At day 0, for the flumes receiving the -Cu treatment 500 mL of the recirculating water were removed and replaced by, 500 mL of an *E. coli* CM2372 suspension in sterilized river water resulting in a final concentration of 10^7^ *E. coli* CM2372 mL^-1^ in the flume. For the +Cu treatment, CuSO_4_ resulting in a final concentration of 500 ppb of Cu^2+^ in the flume was additionally added with the 500 mL of sterilized river water. For the control exclusively, sterilized river water only was pumped into the tank.

### Biofilm sampling

For each destructive sampling of biofilms, an individual glass slide with grown biofilm was randomly removed from each of the three replicate AEUs of each flume. The remaining surrounding water was removed and biofilms were scraped from each individual glass slide inside an individual 50 mL centrifuge tube using a cell scraper (TPP, Trasadingen, Switzerland). One mL of detergent solution (0.9% sterile NaCl solution with 0.05 % of Tween80 (Sigma-Aldrich, St. Louis, MO, USA) was added to support the scrapping process and allow complete detachment of the biofilm from the glass slide. The collected biofilms were then centrifuged at 4000 rpm for 10 minutes and the supernatant was carefully removed by pipetting. The resulting biofilm pellet was weighed and frozen at -20°C until subsequent DNA extraction, carried out using the Qiagen Dneasy PowerSoil Pro Kit, (Qiagen, Hilden, Germany) according to the instruction manual.

For each AEU, an individual glass slide containing biofilm was sampled right after collection from the river to assess the initial diversity of the microbiome (initial time point, T_i_). After the seven-day acclimatization period in the flumes, the biofilms of each AEU were sampled again at experimental day 0 (T_0_) right before applying the respective treatments. Thereafter, biofilm from each AEU were sampled at days 1, 2, 7 and 14 after applying the treatments.

### qPCR based determination of *E. coli* establishment in the biofilm

To determine the success of *E. coli* CM2372 establishment in the biofilm, the absolute abundance of total bacteria, based on the bacterial 16s rRNA gene, and the abundance of *E. coli* CM2372, based on the above described specific sequences, were quantified for all the DNA extracts using Real-time qPCR in a C1000 Touch™ Thermal cycler (Biorad, Hercules, CA, USA) and previously published primer and probe sequences (Table 1). As a qPCR standard the recombinant plasmid pBELX-1 [41], containing both targets, was extracted using the Wizard plus SV Miniprep DNA Purification System (Promega, Madison, WI, USA) according to the manufacturer instructions and linearized by BamHI (Promega) before being purified with the QIAquick PCR purification kit (Qiagen).

**Table 1:**
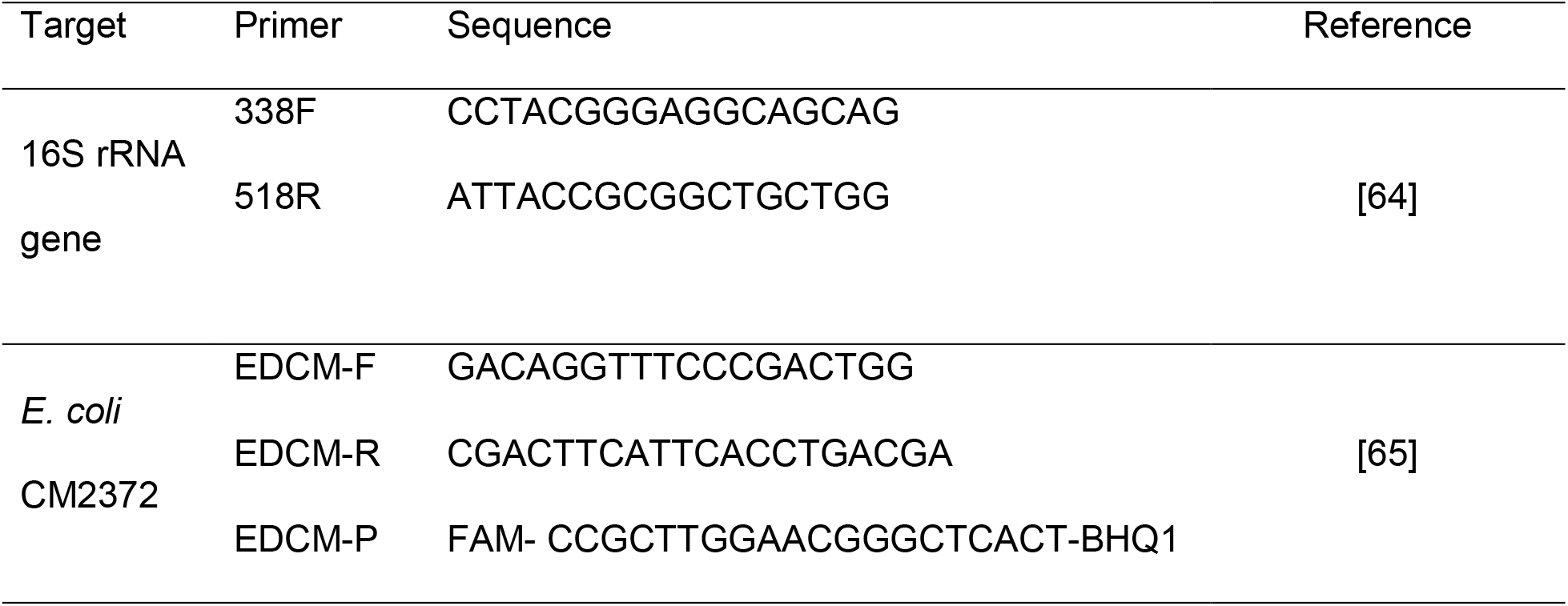
qPCR primers and probes for the detection of *E. coli* CM2372 and the 16S rRNA gene

Reactions were performed in technical triplicates in a MasterCycler RealPlex (Eppendorf, Hamburg, Germany) at a final volume of 20 μL with 10 μL of Luna® Universal qPCR Master Mix (New England Biolabs, Frankfurt, Germany) for the 16S rRNA gene, and 10 μL Luna® Universal Probe qPCR Master Mix (New England Biolabs) for *E. coli* CM2372. Each primer was added at a final concentration of 300 nM, the probe for *E. coli* detection was added at a final concentration of 80 nM, and the reactions were run with 10 ng of DNA extract. The PCR program consisted of initial denaturation at 95 °C for 10 min and 40 cycles of denaturation (95°C; 15 s) and annealing and elongation (60 °C; 1 min). Standard curves for either of the targets were created during every qPCR run, using the above described standard plasmid (containing both targets) with the standard target concentrations ranging from 10^6^ - 10^1^ copies per reaction. Standard curves with amplification efficiency 0.9–1.1 and R^2^ ≥ 0.99 were accepted and melting curve analysis was performed to assess the amplicons’ specificity in the case of 16S rRNA gene quantification. Screening for PCR inhibition was performed by spiking the standard plasmid into the DNA samples. No inhibition was detected in any of the samples. Importantly, no amplification of the *E. coli* target was observed for any of the samples from the control treatment, hence verifying that exclusively the focal *E. coli* strain CM2372, and no unspecific environmental bacteria are detected using this set of primers and probe. The limit of quantification was calculated for each individual qPCR run according to the MIQE guidelines [42]. The absolute abundance of genes was finally expressed as gene copies/cm^2^ on the glass slides and the relative abundance of *E. coli* CM2372 was calculated as the ratio of target copies per copy of the 16S rRNA gene.

### 16S sequencing and analysis of microbial community diversity

The microbial community diversity and structure was profiled using high-throughput amplicon sequencing of the v3-v4 region of the 16S rRNA gene. The DNA extracts were sent to the IKMB Kiel University (Germany) and the 16S rRNA genes were sequenced on an Illumina Novaseq using the primers (v3f: CCTACGGGAGGCAGCAG; v4r: GGACTACHVGGGTWTCTAAT). Sequence analysis was carried out using mothur v.1.47.0 [43] according to the MiSeq SOP [44] as assessed on 03.07.2022. Sequences were classified based on the RDP classifier [45]. Diversity was assessed based on observed OTUs at 97% sequence similarity using the Shannon diversity, Shannon evenness and Chao1 Richness algorithms. Differences in beta diversity between groups of samples was tested using an AMOVA test within the mothur environment [43]. All sequencing data presented in this manuscript have been submitted to the NCBI sequencing read archive under accession number PRJNA903094.

## Results

### Contrasting diversity of the two tested river microbiomes

To understand the invasion dynamics of foreign bacteria into river microbial communities under stress conditions, sites at two different rivers (HAU & HIR) were chosen for the experiment. Natural biofilms were allowed to grow on the glass slides in the AEUs in these rivers, transferred to the laboratory flume system and their initial microbial diversity was characterized. The Shannon diversity index of the first river, HAU, was 6.55 ± 0.22, and significantly higher than that of HIR, 5.53 ± 0.64 (two tailed t-test, p<0.001, df=15) (Fig. 1A). The beta-diversity of HAU and HIR was also significantly different from each other (AMOVA, FS = 20.6069, p<0.001) based on Bray-Curtis dissimilarity (Fig. 1B).

**Figure 1:**
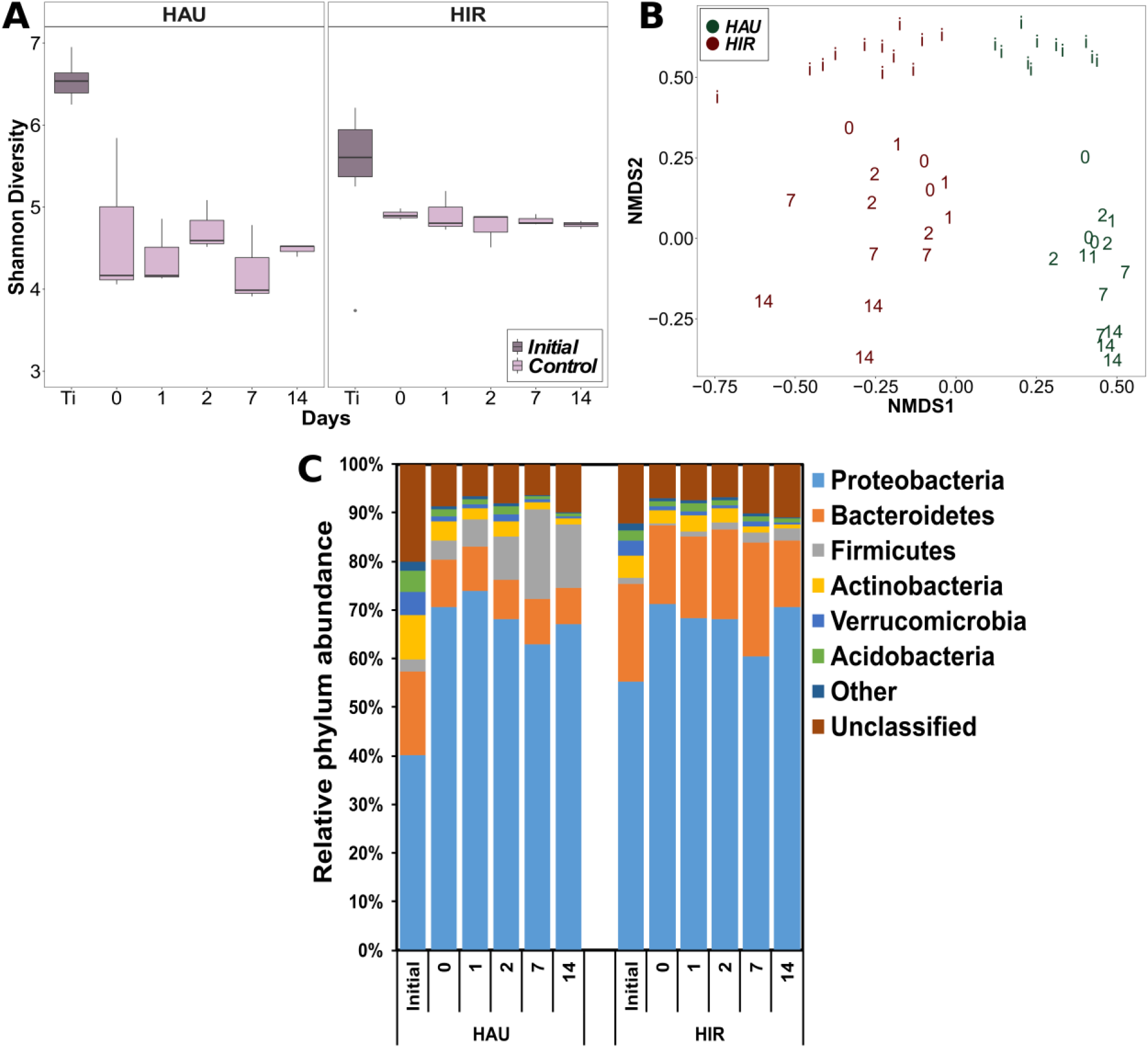
Diversity of river biofilm communities obtained from Hausdorfer Bach (HAU) and Hirschbach (HIR) from the time the biofilm was retrieved from the river (initial = i) and from the control flume without *E. coli* CM2372 nor Cu^2+^ addition. Numbers indicate the day of sampling during the flume experiment. (A) Boxplot of alpha (Shannon) diversity of HAU and HIR. The boxes represent the median with the lower and upper quartile range. Datapoints outside the box are considered outliers. (B) Non-metric multidimensional scaling (NMDS) analysis using Bray-Curtis dissimilarity of beta diversity of the river biofilm communities. (C) Diversity of the microbial biofilm communities on the phylum level (mean relative abundance from replicates is displayed). Only dominant phyla with relative abundance of at least 1% in at least one of the samples are shown. Phyla with lower abundance are grouped as “Other”.

The dominant phylum in both microbial communities was Proteobacteria (40.27 ± 5.02% HAU; 55.08 ± 2.40% HIR), followed by Bacteroidetes (16.86 ± 2.42% HAU; 19.84 ± 1.06% HIR) (Fig. 1C). While on the phylum level, no obvious differences were observed, on the OTU level, the most commonly observed genera in HAU belonged to *Flavobacterium* (2.88 ± 1.41 %), *Acidobacteria* Gp6 (1.35 ± 0.7%) and *Iluminabacter* (*Acidimicrobiaceae*) (1.3 ± 1.35%). Contrary, in HIR they were *Novosphingobium* (*Sphingomonadaceae*) (4.38 ± 1.53%), *Flavobacterium* (3.44 ± 2.53%) and *Emticicia* (*Cytophagaceae*) (1.4 ± 01.23%).

The biofilms were placed inside the flumes for a week to allow the microbial communities to acclimatize to the laboratory conditions before starting the respective treatments. After the acclimatization period, the Shannon diversity of the biofilms decreased from the initial sampling to day 0 for both HAU and HIR (HAU: p<0.05; HIR p<0.05, Mann-Whitney U test; Fig. 1A). The beta diversity of the microbial communities for both HAU and HIR shifted as well, from its initial diversity after the acclimatization period to day 0 (both FS = 20.6069, p<0.05; AMOVA; Fig. 1B). However, the biofilm diversity inside the control flumes remained stable for both rivers from day 0 till day 14. The Shannon diversity index of the microbial community of the HAU control treatment at day 0 was 4.68 ± 1.00 and did not significantly change till day 14 (4.48 ± 0.07; p>0.5, Mann-Whitney U test). Similarly, the diversity for the HIR control at day 0 (4.9 ± 0.07) did not significantly differ from day 14 (4.79 ± 0.05; p>0.05, Mann-Whitney U test) (Fig. 1A). Richness and evenness of the control groups followed similar trends (Fig. SI2 & SI3). This means that subsequent changes in diversity during the experiments would be the consequence of the treatments applied.

Overall, two river microbiomes with originally differing and well characterized diversities were obtained for subsequent invasion experiments. After an initial drop in diversity due to the adaptation of the communities to laboratory conditions, their Shannon diversity remained stable throughout the experiments. This allowed comparing effects across different treatments and communities.

### Initial introduction and subsequent establishment of *E. coli* into the biofilm communities

To understand the invasion dynamics of foreign resistant bacteria into the river microbial communities in the presence and absence of copper-induced stress, the biofilms obtained from HAU and HIR were exposed to a single amendment of *E. coli* CM2372 and monitored for the introduction and establishment success of the focal strain into the biofilm over a period of 14 days. For both rivers, three flumes with different treatments were run: a) only exposed to *E. coli* (-Cu), b) exposed to *E. coli* and Cu^2+^ (+Cu) and c) a control group exposed to neither *E. coli* nor Cu^2+^. During the 14-day period, the absolute amount of biofilm associated bacteria (16S rRNA genes per cm^2^) on each glass slide did not significantly change from those observed at day 0 based on the slope of the Pearson correlation being not significantly different from /zero0 (HAU: p>0.05; HIR: p>0.05; Fig. 2) for either of the tested rivers or treatments.

**Figure 2:**
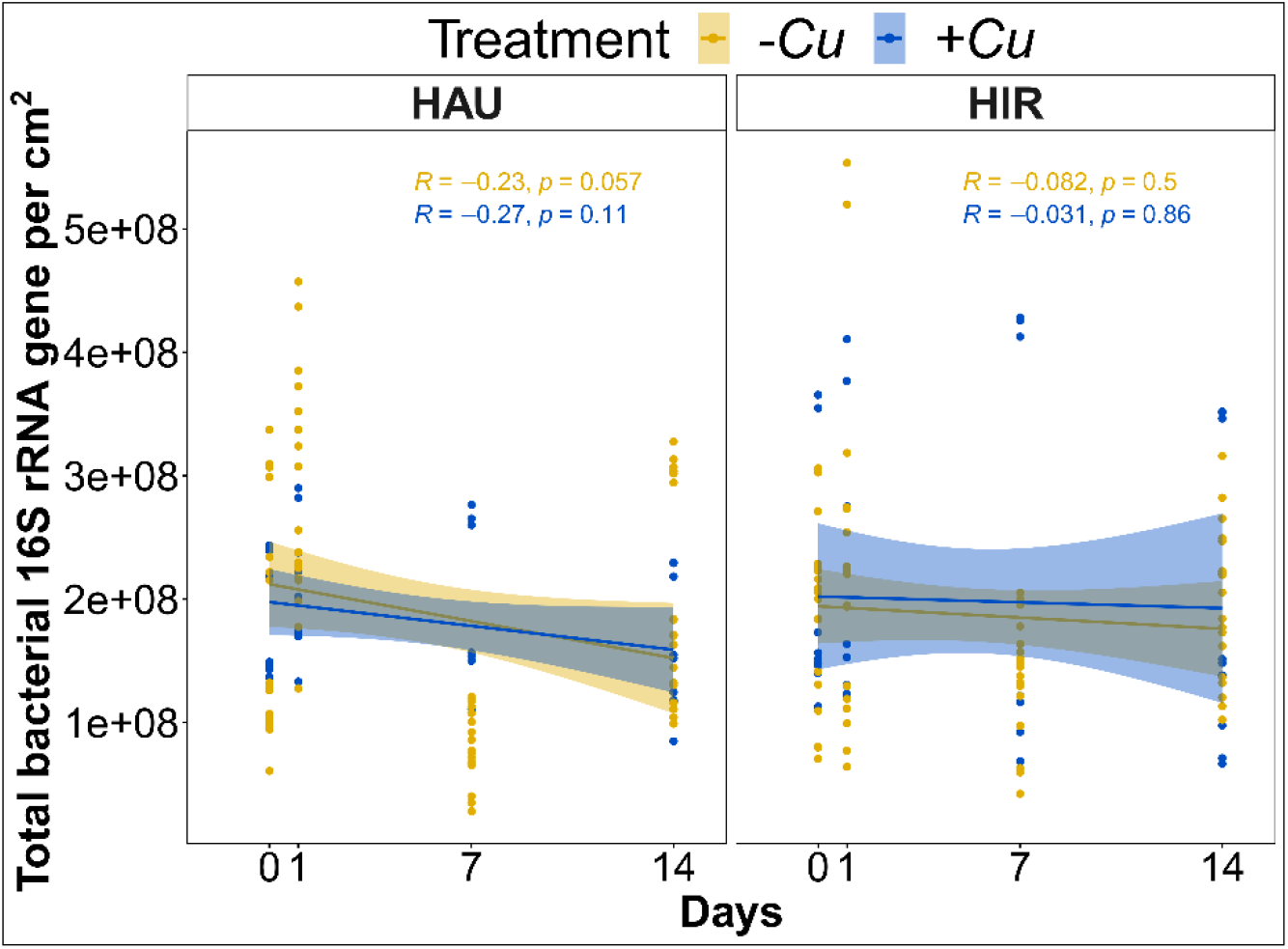
Total bacterial 16S rRNA gene abundance in the biofilms on the glass slides per cm^2^ in the flumes did not significantly change over time for any of the rivers Hausdorfer Bach (HAU) and Hirschbach (HIR), or any of the treatments with (+Cu) or without (-Cu) the addition of Cu^2+^.

Furthermore, there was no significant difference between the absolute bacterial abundance when comparing the -Cu and +Cu treatment for either of the rivers (HAU: p>0.05; HIR: p>0.05; Fig. 2). As the total amount of 16S rRNA genes did not change over time or treatment, changes in the relative abundance of *E. coli* CM2372 per total 16S rRNA gene do represent changes in its absolute abundance, hence success of invasion and establishment.

On day 1, all biofilms that were exposed to *E. coli* CM2372 displayed a measurable amount of the focal *E. coli* strain that got introduced to the biofilm. In HAU -Cu, the *E. coli* invader strain reached a relative abundance of 3.98 ± 2.39 × 10^−3^, while for HAU +Cu, the relative abundance of *E. coli* CM2372 associated to biofilms was slightly lower with 2.21 ± 0.46 × 10^−3^ (Fig. 3A). The relative abundance for *E. coli* in HIR -Cu biofilms was 2.87 ± 0.94 × 10^−3^ and slightly higher for HIR +Cu with 3.73 ± 0.74 × 10^−3^. There was no clear trend that could be associated to any treatment effect as the comparisons of relative abundances of *E. coli* CM2372 associated biofilms revealed no significant differences for either HAU (w=54, p=1.00, Mann-Whitney U test) or HIR (w=37, p=0.13878, Mann-Whitney U test) (Fig. 3A). Hence, initially *E. coli* can successfully be introduced to the biofilm communities independent of the imposed stress conditions or the origin of the biofilm.

**Figure 3:**
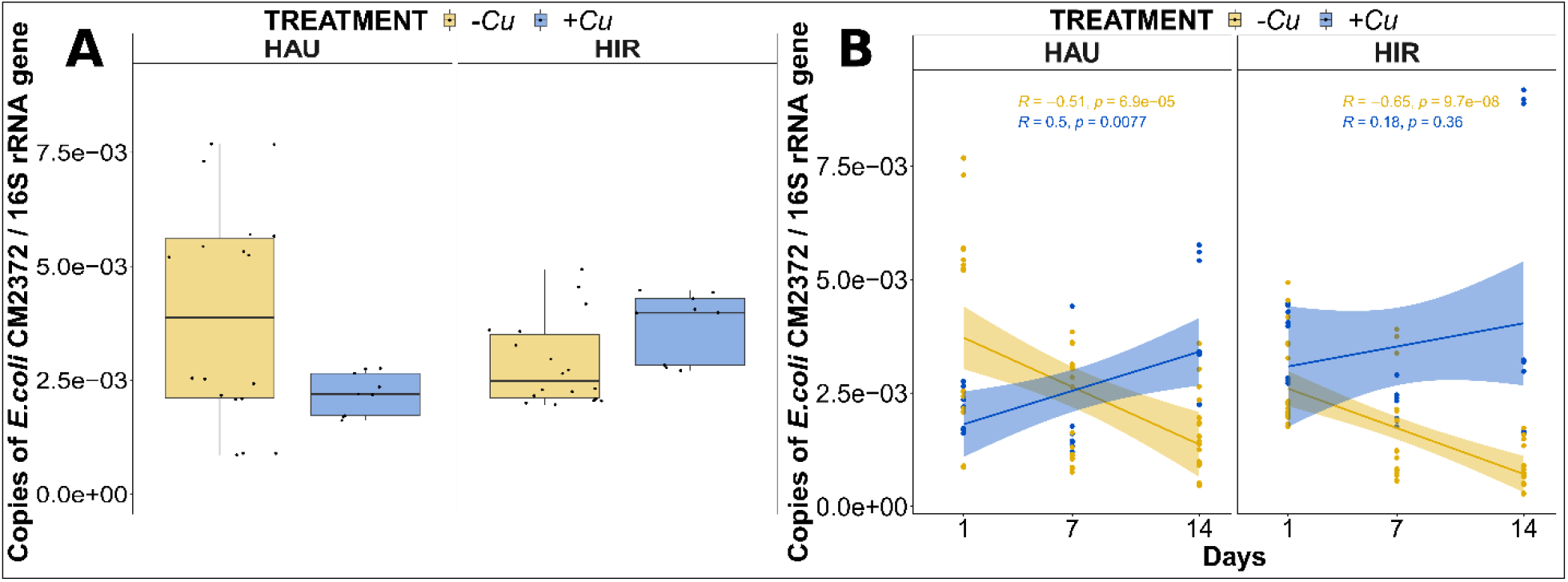
Invasion and establishment of *E. coli* CM2372 into the river microbial biofilm communities. The relative abundance of *E. coli* in log scale for both river microbial communities from Hausdorfer Bach (HAU) and Hirschbach (HIR) with (+Cu) or without copper (-Cu). (A) Relative *E. coli* CM2372 abundance in log scale for day 1, the day after inoculation of *E. coli* strain representing the initial introduction into the biofilms. (B) Linear regression for relative *E. coli* CM2372 abundance over time in the biofilm.

However, following the initial step of introduction to the biofilm, the relative abundance of *E. coli* CM2372 decreased gradually after day 1 in the absence of Cu^2+^ for both rivers, with significantly negative slopes observed based on Pearson correlation (p<0.001, Fig. 3B). After 14 days, the relative abundance of *E. coli* CM2372 was reduced by 60% for HAU -Cu to 1.6 ± 0.89 × 10^−3^. Similarly, for HIR -Cu, the relative abundance was reduced by 70% to 0.81 ± 0.45 × 10^−3^. Meanwhile, for the +Cu treatment in both HAU and HIR, a successful establishment of *E. coli* into the community post initial introduction could be observed. The relative abundance of *E. coli* stayed relatively stable for HIR, reaching 4.6 ± 3.4 × 10^−3^ at day 14 with a slightly positive slope that was not significantly different from 0 (p=0.36, Pearson correlation). Further, it increased significantly for HAU by 41.05% after 14 days to 3.8 ± 1.5 × 10^−3^ with the slope being significantly positive (p<0.001, Pearson correlation).

To confirm if the relative abundance of *E. coli* CM2372 as a dependent variable can be explained using the two independent variables time (time) and presence or absence of copper (Cu), we set up a linear model (Eq. 1) describing their individual contribution and the contribution of their interaction:

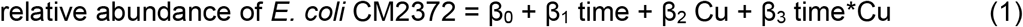

The model was run for the datasets from each river individually. Time on its own had a significantly negative effect (β_1_<0) on the abundance of *E. coli* CM2372 in biofilms from both rivers (HAU: p<0.001, HIR: p<0.001), mirroring the observation that in the absence of copper the focal *E. coli* strain decreased in abundance. Copper on its own (β_2_) had no significant effect on the initial relative abundance of *E. coli* CM2372 as described above. However, the interaction of copper and time had throughout a significantly positive effect (β_3_>0) on the relative abundance of *E. coli* CM2372 (HAU: p<0.001, HIR: p=0.001). In both cases, the positive effect of the copper x time interaction was stronger than the negative effect of time on its own (|β_3_| > |β_1_|), confirming that the presence of copper as a stressor resulted in the observed increase in relative *E. coli* CM2372 abundance and hence contributed to the elevated invasion success.

In summary, despite the initial successful introduction of *E. coli* CM2372 into the biofilms, the invader did not establish itself into the biofilm communities in the absence of stress. However, under stress induced by copper, establishment and potentially proliferation of the invading strain became possible.

### Loss in microbial diversity under copper stress correlates with increased establishment of the invader strain

As contrasting persistence dynamics of *E. coli* CM2372 into the biofilm communities in the presence and absence of stress were observed, we investigated whether stress induced shifts in microbial diversity and structure of the biofilm communities could provide an explanation for the observed dynamics. The Shannon diversity of the microbial communities in the -Cu group was throughout not significant difference from the control group for both HAU (p=0.54, Kruskal-Wallis test) and HIR (p=0.42, Kruskal-Wallis test) (Fig. 4A). Further, no significant changes with time could be observed. Richness and evenness of the -Cu group and control group followed similar trends (Fig. SI2 & SI3). This indicates that there was no direct effect of *E. coli* CM2372 exposure on the diversity of the microbial community.

**Figure 4:**
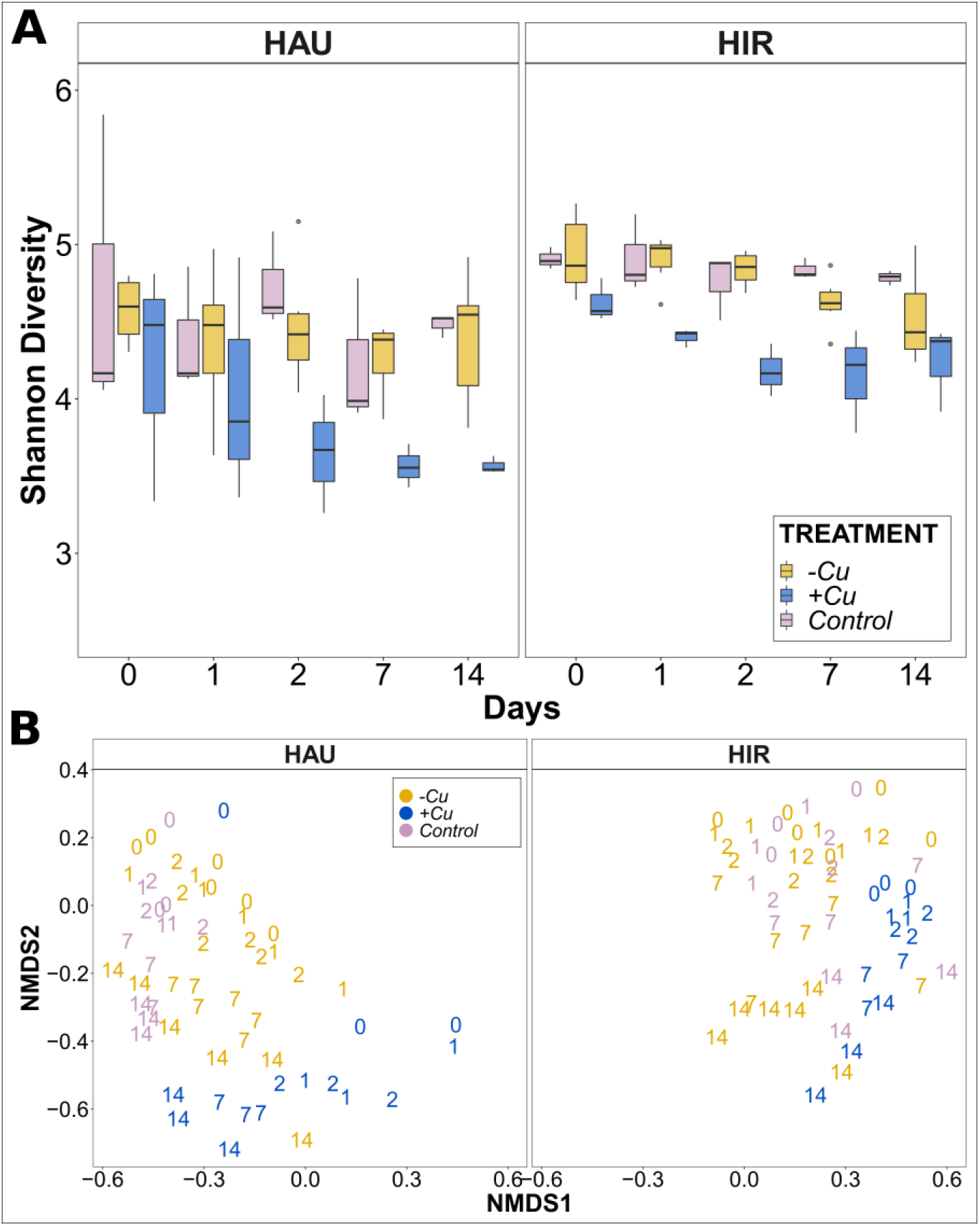
Diversity of the microbial biofilm communities for Hausdorfer Bach (HAU) and Hirschbach (HIR) and the three different treatments of *E. coli* CM2372 with (+Cu) and without (-Cu) the addition of Cu^2+^ inside the flumes over time. Numbers are representing the day of sampling. (A) Alpha diversity: Box plots showing Shannon diversity with the boxes representing the median with the lower and upper quartile range. Datapoints outside the boxes are considered outliers. (B) Beta diversity based on non-metric multidimensional scaling (NMDS) ordinations using Bray-Curtis dissimilarity.

At day 0, the Shannon diversity index of the HAU +Cu treatment was 4.2 ± 0.77 which at day 14 decreased to 3.57 ± 0.05, substantiating a significant shift in the diversity between -Cu and +Cu over time (p<0.001, Kruskal-Wallis test, Fig. 4A). Similar trends were observed for the second river, HIR, with a significant difference in the diversity between -Cu and +Cu after 14 days (p<0.001, Kruskal-Wallis test, Fig. 4A).

The beta diversity for each of the two rivers equally showed dissimilarities between different treatment groups (Fig. 4B). For both rivers, the biofilms that received the control and the –Cu treatment again clustered together. The beta diversity of the +Cu treatment group clustered away from the other two treatment groups for both HAU and HIR (AMOVA, p<0.001) (Fig. 4B).

Hence, it is evident that the microbial community diversity is disturbed and the community structure is altered under copper stress. This loss in diversity under stress is directly correlated with the observed increase in establishment success of the *E. coli* strain, hence verifying our initial hypothesis.

## Discussion

We here demonstrated that, during the exposure to stressors, a focal invader strain established and increased its abundance within natural river biofilms. The observed dynamics display all signs of a successful biological invasion event consisting of the original introduction, an establishment step and a putative proliferation step. Noteworthy, the increased establishment success of the invader coincided with a loss of microbial community diversity under stress conditions. Contrary, in the unstressed control biofilm of higher diversity no establishment or proliferation of the invader were observed, hence the invasion process can be deemed unsuccessful.

Using the example of a model resistant *E. coli* strain invading natural river microbial biofilms under Cu^2+^ stress, we observed that initially, the invader could attach to the biofilms independent of the presence of stress or the biofilm origin. This was according to invasion theory, as the initial introduction during an invasion process is primarily relying on stochastic processes [26, 29]. The initial introduction success depends mainly on the quantity of invaders present, also known as the propagule pressure as well as the attachment quality of the invader [46, 47]. The absolute numbers of invaders as well as indigenous biofilm community members were consistent between treatments and rivers of origin. Consequently, this propagule pressure can also be assumed as constant in the experiments. As throughout the identical invader strain is used, the quality of the invader attachment ability can similarly be assumed as equal. The observed stochasticity of the initial attachment process, independent of copper induced stress, is hence a logical consequence.

Such initial introduction events of invaders of anthropogenic origin can be assumed as ubiquitous in the environment. High numbers of resistant bacteria are consistently released through wastewater effluents into rivers. For example, on average 10^11^ cefotaxime resistant coliform bacteria get daily released from large, urban WWTPs into the receiving rivers based on a global survey [34]. Compared to the quantity of invading *E. coli* used in the experiments here, the final concentration of coliform bacteria in receiving water bodies in industrialized countries are usually slightly lower [48]. However, rivers in places with poor or no WWTP infrastructure consistently receive sewage without any prior or only partial treatment and regularly exceed concentrations of 10^6^ coliforms mL^-1^ [49–52]. Moreover, the release of stressors co-introduced with the bacterial contaminants is regularly higher in those places, which makes the here presented results highly environmentally relevant. Still, as demonstrated above, the initial introduction process is independent of these stressors. However, it provides the basis for the next step during the invasion process, the subsequent establishment in the natural river biofilm communities, for which we demonstrate a clear effect of co-released microbial stressors.

Once the initial attachment to the biofilm community, hence the introduction of the invader, is completed, the internal resistance of the microbial community towards invasion, e.g. biotic barriers, have to be overcome to lead to the successful establishment of the invader [26, 29]. Changes in the community composition and diversity can result in opening up available niches that become accessible to the invading bacteria [28, 30, 35, 53, 54]. A community’s invasion resistance is related to the available niches for invader establishment: here we suspect that more diverse communities provide less available niche spaces due to a higher degree of different bacteria interacting together for the resources available. This is commonly known as the “diversity-invasion effect” [26, 29]. In our experiments, the microbial communities in their original state (in the absence of stress) can better resist the invasion by *E. coli* CM2372 as biotic barriers towards invasion, due to less available niche space, cannot be overcome. This results in the invading *E. coli* population perishing from the microbial community after initial introduction. Moreover, it implies that highly diverse microbial communities can serve as a first line of defense towards invasion by resistant bacteria of anthropogenic origin in the environment. In contrast, invading *E. coli* could thrive and establish a stable population in the newly invaded biofilms when there was a loss in the river microbial communities’ diversity under copper stress. Similar effects of environmental stresses on the diversity of the microbial communities have regularly been reported for soils and rivers [29, 31–33, 55, 56]. The observed loss in diversity coincides with the opening of niches within the community, resulting in gaps in the biotic barrier towards invasion that can be exploited by the invaders for their establishment, here demonstrated for *E. coli*. Indeed, *E. coli* have been shown to possess high levels of survival and persistence in diverse environment and even have their own environmental life cycles [57–59], a clear indication that at least transiently successful invasion events of resistant *E. coli* into the river microbiome will regularly take place. Within this context, it is further interesting to note that, while metals have long been perceived mainly as co-selective agents for ARGs in the environment [7], they may additionally be simple community structure disruptors that allow for a more successful invasion of bacteria originating from anthropic environments where the selective enrichment of AMR is already well documented.

In the context of wastewater effluents entering rivers, this means that the invasion of resistant bacteria is supported by the constant co-release of stressors in the effluents. This in turn leads to increased success in their establishment, with the invading resistant bacteria becoming either transient or permanent members of the community, hence increasing their abundance. This is concerning as even a prolonged residence time can favor the spread of ARGs via horizontal gene transfer from the invading foreign bacteria to the indigenous communities, thus leading to an even higher persistence of ARGs [41]. In the presence of environmental stressors, such horizontal gene transfer rates are regularly elevated [21, 22, 60, 61]. Furthermore, even if the invasion is only transiently successful and no horizontal transfer has taken place, the invading bacteria can leave a permanent dent in the original microbial community and niche structure, by *e*.*g*. knocking out key species of the community [29, 62]. This loss in diversity resilience, in turn, leaves the community vulnerable to following consecutive invasion events [26, 29, 62]. Consequently, to combat the spread of AMR through wastewater effluents into river microbiomes, “an essential battlefront in the war on antimicrobial resistance” [63], both permanent as well as transiently successful invasion events of resistant bacteria need to be limited.

We here demonstrated that microbial community resistance towards invasion by foreign resistant bacteria is directly affected by disturbances through environmental stressors. The microbial community diversity shifts under stress conditions, which in turn increases its susceptibility towards invasion by resistant bacteria. Therefore, it is important to reevaluate not only limits regarding the release of resistant bacteria but to consider it in context of the co-released environmental stressors and their consequences on the structure of the downstream microbial communities.

## Acknowledgements

The authors thank Lena Winter, Emily Fischer and Steffen Kunze for help in sampling and setting up and running the flume experiments. We thank David Kneis and Johannes Feldbauer for support with the model. We further thank the members of the ANTIVERSA consortium for their contributions in designing the flume setup.

## Funding

This work was supported by the ANTIVERSA project (BiodivERsa2018-A-452) here funded by the Bundesministerium für Bildung, und Forschung of Germany under grant number 01LC1904A and the Agence Nationale de la Recherche under grant number ANR-19-EBI3-0005-04. KB was supported through a DAAD scholarship in the program Research Grants - Bi-nationally Supervised Doctoral Degrees/Cotutelle, 2021/22 (57552338). GS was supported through a Faculty Initiation Grant (FIG/100762) and Department of Science and Technology under grant number DST/TM/INDO-UK/2K17/46. Responsibility for the information and views expressed in the manuscript lies entirely with the authors.

## Competing Interests

The authors declare no competing interests.

## Notes

### Competing Interest Statement

The authors have declared no competing interest.

## References

1. O’Neill J. Tackling drug-resistant infections globally: Report and recommendations. The review on antimicrobial resistance chaired by Jim O’Neill. 2016.

2. Hernando-Amado S, Coque TM, Baquero F, Martínez JL. Defining and combating antibiotic resistance from One Health and Global Health perspectives. Nature Microbiology 2019 4:9 2019; 4: 1432–1442.

3. McEwen SA, Collignon PJ. Antimicrobial Resistance: a One Health Perspective. Microbiol Spectr 2018; 6.

4. Karkman A, Pärnänen K, Larsson DGJ. Fecal pollution can explain antibiotic resistance gene abundances in anthropogenically impacted environments. Nat Commun 2019; 10.

5. Pruden A, Arabi M, Storteboom HN. Correlation Between Upstream Human Activities and Riverine Antibiotic Resistance Genes. 2012.

6. Berendonk TU, Manaia CM, Merlin C, Fatta-Kassinos D, Cytryn E, Walsh F, et al. Tackling antibiotic resistance: The environmental framework. Nat Rev Microbiol 2015; 13: 310–317.

7. Pal C, Bengtsson-Palme J, Kristiansson E, Larsson DGJ. Co-occurrence of resistance genes to antibiotics, biocides and metals reveals novel insights into their co-selection potential. BMC Genomics 2015; 16.

8. Marathe NP, Pal C, Gaikwad SS, Jonsson V, Kristiansson E, Larsson DGJ. Untreated urban waste contaminates Indian river sediments with resistance genes to last resort antibiotics. Water Res 2017; 124: 388–397.

9. Pal C, Asiani K, Arya S, Rensing C, Stekel DJ, Larsson DGJ, et al. Metal Resistance and Its Association With Antibiotic Resistance. Adv Microb Physiol 2017; 70: 261–313.

10. Heß S, Berendonk TU, Kneis D. Antibiotic resistant bacteria and resistance genes in the bottom sediment of a small stream and the potential impact of remobilization. FEMS Microbiol Ecol 2018; 94.

11. Huijbers PMC, Blaak H, de Jong MCM, Graat EAM, Vandenbroucke-Grauls CMJE, de Roda Husman AM. Role of the Environment in the Transmission of Antimicrobial Resistance to Humans: A Review. Environ Sci Technol 2015; 49: 11993–12004.

12. Li A-D, Li L-G, Zhang T. Exploring antibiotic resistance genes and metal resistance genes in plasmid metagenomes from wastewater treatment plants. Front Microbiol 2015; 6.

13. Munir M, Wong K, Xagoraraki I. Release of antibiotic resistant bacteria and genes in the effluent and biosolids of five wastewater utilities in Michigan. Water Res 2011; 45: 681–693.

14. Rizzo L, Manaia C, Merlin C, Schwartz T, Dagot C, Ploy MC, et al. Science of the Total Environment Urban wastewater treatment plants as hotspots for antibiotic resistant bacteria and genes spread into the environment : A review. Science of the Total Environment, The 2013; 447: 345–360.

15. Malagón-Rojas JN, Parra Barrera EL, Lagos L. From environment to clinic: the role of pesticides in antimicrobial resistance. Revista Panamericana de Salud Pública 2020; 44.

16. Ramakrishnan B, Venkateswarlu K, Sethunathan N, Megharaj M. Local applications but global implications: Can pesticides drive microorganisms to develop antimicrobial resistance? Science of The Total Environment 2019; 654: 177–189.

17. aus der Beek T, Weber FA, Bergmann A, Hickmann S, Ebert I, Hein A, et al. Pharmaceuticals in the environment—Global occurrences and perspectives. Environ Toxicol Chem 2016; 35: 823–835.

18. Larsson DGJ, de Pedro C, Paxeus N. Effluent from drug manufactures contains extremely high levels of pharmaceuticals. J Hazard Mater 2007; 148: 751–755.

19. Reddy B, Dubey SK. River Ganges water as reservoir of microbes with antibiotic and metal ion resistance genes : High throughput metagenomic approach *. Environmental Pollution 2019; 246: 443–451.

20. Stanton IC, Tipper HJ, Chau K, Klümper U, Subirats J, Murray AK. Does Environmental Exposure to Pharmaceutical and Personal Care Product Residues Result in the Selection of Antimicrobial-Resistant Microorganisms, and is This Important in Terms of Human Health Outcomes? Environ Toxicol Chem 2022.

21. Klümper U, Dechesne A, Riber L, Brandt KK, Gülay A, Sørensen SJ, et al. Metal stressors consistently modulate bacterial conjugal plasmid uptake potential in a phylogenetically conserved manner. The ISME Journal 2017 11:1 2016; 11: 152–165.

22. Klümper U, Riber L, Dechesne A, Sannazzarro A, Hansen LH, Sørensen SJ, et al. Broad host range plasmids can invade an unexpectedly diverse fraction of a soil bacterial community. ISME Journal 2015; 9: 934–945.

23. Lu J, Wang Y, Li J, Mao L, Nguyen SH, Duarte T, et al. Triclosan at environmentally relevant concentrations promotes horizontal transfer of multidrug resistance genes within and across bacterial genera. Environ Int 2018; 121: 1217–1226.

24. Huijbers PMC, Flach CF, Larsson DGJ. A conceptual framework for the environmental surveillance of antibiotics and antibiotic resistance. Environ Int 2019; 130.

25. Karkman A, Pärnänen K, Larsson DGJ. Fecal pollution can explain antibiotic resistance gene abundances in anthropogenically impacted environments. Nat Commun 2019; 10: 80.

26. Mallon CA, van Elsas JD, Salles JF. Microbial Invasions: The Process, Patterns, and Mechanisms. Trends Microbiol 2015; 23: 719–729.

27. Kinnunen M, Dechesne A, Proctor C, Hammes F, Johnson D, Quintela-Baluja M, et al. A conceptual framework for invasion in microbial communities. The ISME Journal 2016 10:12 2016; 10: 2773–2779.

28. Finke DL, Snyder WE. Niche partitioning increases resource exploitation by diverse communities. Science (1979) 2008; 321: 1488–1490.

29. Mallon CA, le Roux X, van Doorn GS, Dini-Andreote F, Poly F, Salles JF. The impact of failure: unsuccessful bacterial invasions steer the soil microbial community away from the invader’s niche. The ISME Journal 2018 12:3 2018; 12: 728–741.

30. de Roy K, Marzorati M, Negroni A, Thas O, Balloi A, Fava F, et al. Environmental conditions and community evenness determine the outcome of biological invasion. Nature Communications 2013 4:1 2013; 4: 1–5.

31. Zhao J, Zhao X, Chao L, Zhang W, You T, Zhang J. Diversity change of microbial communities responding to zinc and arsenic pollution in a river of northeastern China. J Zhejiang Univ Sci B 2014; 15: 670–680.

32. Wu H, Li Y, Zhang W, Wang C, Wang P, Niu L, et al. Bacterial community composition and function shift with the aggravation of water quality in a heavily polluted river. J Environ Manage 2019; 237: 433–441.

33. García-Armisen T, İnceoğlu Ö, Ouattara NK, Anzil A, Verbanck MA, Brion N, et al. Seasonal Variations and Resilience of Bacterial Communities in a Sewage Polluted Urban River. PLoS One 2014; 9: e92579.

34. Marano RBM, Fernandes T, Manaia CM, Nunes O, Morrison D, Berendonk TU, et al. A global multinational survey of cefotaxime-resistant coliforms in urban wastewater treatment plants. Environ Int 2020; 144: 106035.

35. Xing J, Chen M, Deng X, Chen J, Jiang P, Qin H. Resilience of soil microbial metabolic functions to temporary E. coli invasion. 2022.

36. Ishii S, Ksoll WB, Hicks RE, Sadowsky MJ. Presence and growth of naturalized Escherichia coli in temperate soils from lake superior watersheds. Appl Environ Microbiol 2006; 72: 612–621.

37. Song J, Klümper U, Riber L, Dechesne A, Smets BF, Sørensen SJ, et al. A converging subset of soil bacterial taxa is permissive to the IncP-1 plasmid pKJK5 across a range of soil copper contamination. FEMS Microbiol Ecol 2020; 96: 1–10.

38. Seiler C, Berendonk TU. Heavy metal driven co-selection of antibiotic resistance in soil and water bodies impacted by agriculture and aquaculture. Front Microbiol 2012; 3: 399.

39. Merlin C, Gardiner G, Durand S, Masters M. The Escherichia coli metD locus encodes an ABC transporter which includes Abc (MetN), YaeE (MetI), and YaeC (MetQ). J Bacteriol 2002; 184: 5513–5517.

40. Smalla K, Heuer H, Gotz A, Niemeyer D, Krogerrecklenfort E, Tietze E. Exogenous isolation of antibiotic resistance plasmids from piggery manure slurries reveals a high prevalence and diversity of IncQ-like plasmids. Appl Environ Microbiol 2000; 66: 4854–4862.

41. Bellanger X, Guilloteau H, Bonot S, Merlin C. Demonstrating plasmid-based horizontal gene transfer in complex environmental matrices: A practical approach for a critical review. Science of The Total Environment 2014; 493: 872–882.

42. Bustin SA, Benes V, Garson JA, Hellemans J, Huggett J, Kubista M, et al. The MIQE guidelines: Minimum information for publication of quantitative real-time PCR experiments. Clin Chem 2009; 55: 611–622.

43. Schloss PD, Westcott SL, Ryabin T, Hall JR, Hartmann M, Hollister EB, et al. Introducing mothur: open-source, platform-independent, community-supported software for describing and comparing microbial communities. Appl Environ Microbiol 2009; 75: 7537–7541.

44. Kozich JJ, Westcott SL, Baxter NT, Highlander SK, Schloss PD. Development of a dual-index sequencing strategy and curation pipeline for analyzing amplicon sequence data on the MiSeq Illumina sequencing platform. Appl Environ Microbiol 2013; 79: 5112–5120.

45. Wang Q, Garrity GM, Tiedje JM, Cole JR. Naïve Bayesian classifier for rapid assignment of rRNA sequences into the new bacterial taxonomy. Appl Environ Microbiol 2007; 73: 5261–5267.

46. Acosta F, Zamor RM, Najar FZ, Roe BA, Hambright KD. Dynamics of an experimental microbial invasion. Proc Natl Acad Sci U S A 2015; 112: 11594–11599.

47. Kinnunen M, Dechesne A, Albrechtsen HJ, Smets BF. Stochastic processes govern invasion success in microbial communities when the invader is phylogenetically close to resident bacteria. The ISME Journal 2018 12:11 2018; 12: 2748–2756.

48. Cacace D, Fatta-Kassinos D, Manaia CM, Cytryn E, Kreuzinger N, Rizzo L, et al. Antibiotic resistance genes in treated wastewater and in the receiving water bodies: A pan-European survey of urban settings. Water Res 2019; 162: 320–330.

49. Narain S. Sanitation for all. Nature. 2012., 486: 185

50. Mallapur C. 70% Of Urban India’s Sewage Is Untreated. India Spend. 2019., 1–7

51. Williams M, Kookana RS, Mehta A, Yadav SK, Tailor BL, Maheshwari B. Emerging contaminants in a river receiving untreated wastewater from an Indian urban centre. Science of the Total Environment 2019; 647: 1256–1265.

52. Marathe NP, Pal C, Gaikwad SS, Jonsson V, Kristiansson E, Larsson DGJ. Untreated urban waste contaminates Indian river sediments with resistance genes to last resort antibiotics. Water Res 2017; 124: 388–397.

53. Altman S, Whitlatch RB. Effects of small-scale disturbance on invasion success in marine communities. J Exp Mar Biol Ecol 2007; 342: 15–29.

54. Liu X, le Roux X, Salles JF. The legacy of microbial inoculants in agroecosystems and potential for tackling climate change challenges. iScience 2022; 25: 103821.

55. Guo X pan, Yang Y, Lu D pei, Niu Z shun, Feng J nan, Chen Yru, et al. Biofilms as a sink for antibiotic resistance genes (ARGs) in the Yangtze Estuary. Water Res 2018; 129: 277–286.

56. Gao T, Wang X, Liu Y, Wang H, Zuo M, He Y, et al. Characteristics and diversity of microbial communities in lead–zinc tailings under heavy metal stress in north-west China. Lett Appl Microbiol 2022; 74: 277–287.

57. Devane ML, Moriarty E, Weaver L, Cookson A, Gilpin B. Fecal indicator bacteria from environmental sources; strategies for identification to improve water quality monitoring. Water Res 2020; 185: 116204.

58. van Elsas JD, Semenov A v., Costa R, Trevors JT. Survival of Escherichia coli in the environment: fundamental and public health aspects. The ISME Journal 2011 5:2 2010; 5: 173–183.

59. Aijuka M, Buys EM. Persistence of foodborne diarrheagenic Escherichia coli in the agricultural and food production environment: Implications for food safety and public health. Food Microbiol 2019; 82: 363–370.

60. Xing J, Jia X, Wang H, Ma B, Falcão Salles J, Xu J. The legacy of bacterial invasions on soil native communities. Environ Microbiol 2021; 23: 669–681.

61. Wang Y, Yu Z, Ding P, Lu J, Klümper U, Murray AK, et al. Non-antibiotic pharmaceuticals promote conjugative plasmid transfer at a community-wide level. Microbiome 2022; 10: 1–15.

62. Coyte KZ, Schluter J, Foster KR. The ecology of the microbiome: Networks, competition, and stability. Science (1979) 2015; 350: 663–666.

63. Bürgmann H, Frigon D, Gaze WH, Manaia CM, Pruden A, Singer AC, et al. Water and sanitation: an essential battlefront in the war on antimicrobial resistance. FEMS Microbiol Ecol 2018; 94: 101.

64. Muyzer G, de Waal EC, Uitterlinden AG. Profiling of complex microbial populations by denaturing gradient gel electrophoresis analysis of polymerase chain reaction-amplified genes coding for 16S rRNA. Appl Environ Microbiol 1993; 59: 695–700.

65. Sagrillo C, Changey F, Bellanger X. Bacteriophages vehiculate a high amount of antibiotic resistance determinants of bacterial origin in the Orne River ecosystem. Environ Microbiol 2022; 24: 4317–4328.

