## Supplementary Information for "Exposure to environmental stress decreases the resistance of river microbial communities towards invasion by antimicrobial resistant bacteria"

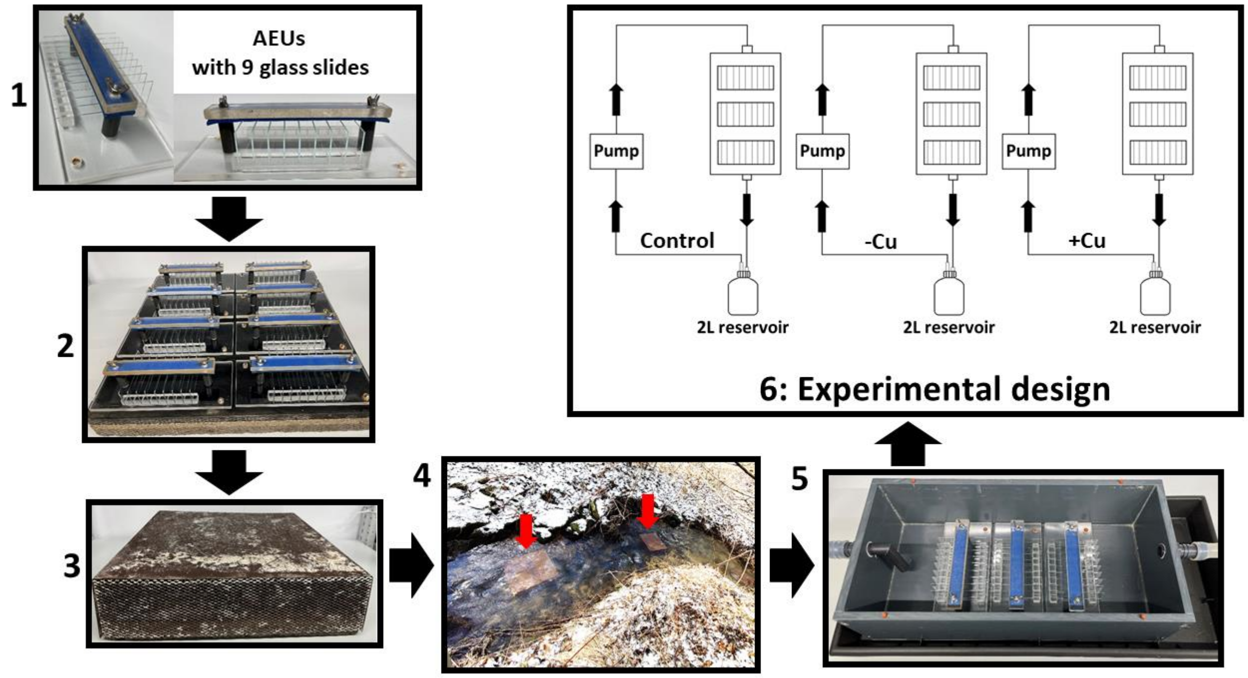


**SI Figure 1:** Schematic workflow of the flume experiments. 1) Biofilms were grown on microscope glass slides of 76 x 26 mm size, with 9 glass slides each fixed in artificial exposure units (AEUs) constructed from Plexiglas. 2) 8 AEUs were screwed onto a concrete plate and 3) covered with a stainless steel cover providing shade and protection with a metal mesh at the front and the back allowing river water to stream through the AEUs. 4) The AEUs were immerged in the rivers Hausdorfer Bach (HAU) and Hirschbach (HIR) for one month to allow biofilm growth. 5) The AEUs containing the glass slides with grown biofilms were then transferred to laboratory artificial flume systems (40 cm long, 19.1 cm width, 10 cm height), made of polypropylene, and equipped with an in- and an outflow pipe at either end. Each flume was connected to a 2L reservoir connected to a recirculation pump and filled with filter sterilized river water constantly recirculated with a pump. 6) Three replicate AEUs containing glass slides covered with river biofilms were placed in the middle of each flume and exposed to three different types of treatments for a period of 14 days: a) solely exposed to E. coli CM2372 (-Cu), b) exposed to E. coli CM2372 and Cu2+ (+Cu) and c) a control group was run without exposure to either E. coli CM2372 or Cu2+. Glass slides containing biofilms were then destructively sampled to monitor the invasion success of E. coli CM2372 into the biofilm over time.

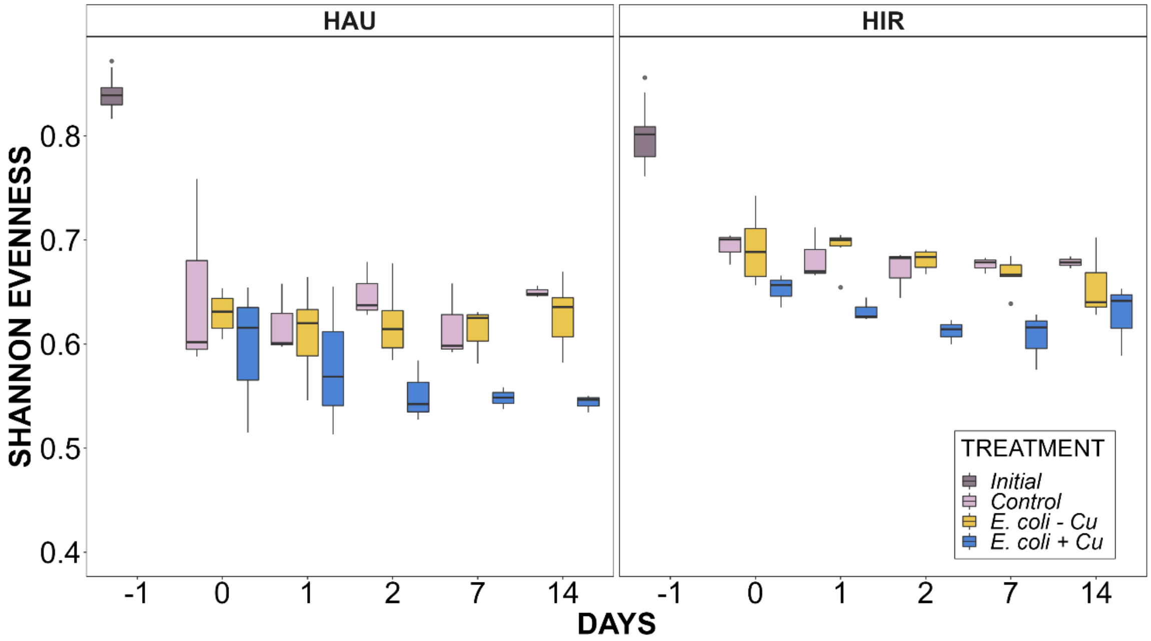


**SI Figure 2:** Shannon evenness of the microbial biofilm communities for Hausdorfer Bach (HAU) and Hirschbach (HIR) and the three different treatments of *E. coli* CM2372 with (+Cu) and without (-Cu) the addition of $Cu^{2+}$ inside the flumes over time. The numbers represent the day of sampling along with the initial Shannon evenness of the community before the experiment started. Box plots with the boxes representing the median with the lower and upper quartile range. Data points outside the boxes are considered outliers.

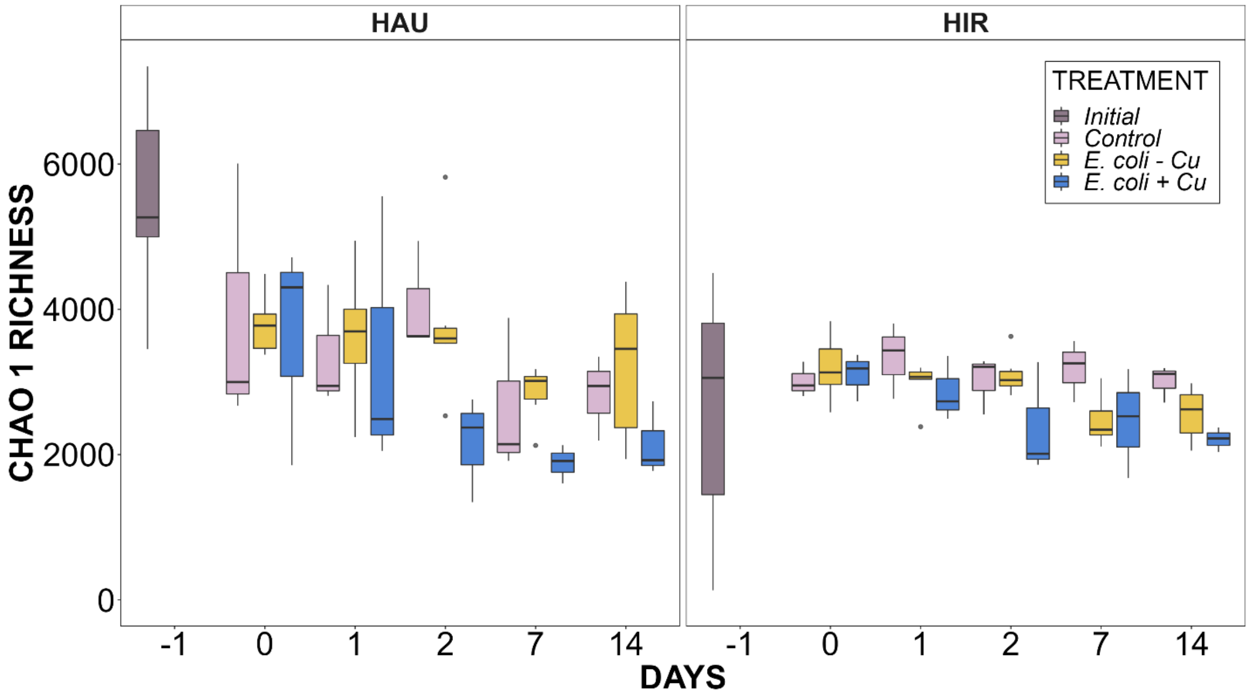


**SI Figure 3:** Chao1 richness of the microbial biofilm communities for Hausdorfer Bach (HAU) and Hirschbach (HIR) and the three different treatments of *E. coli* CM2372 with (+Cu) and without (-Cu) the addition of $Cu^{2+}$ inside the flumes over time. The numbers represent the day of sampling along with the initial Shannon evenness of the community before the experiment started. Box plots with the boxes representing the median with the lower and upper quartile range. Data points outside the boxes are considered outliers.
